# Brain criticality predicts individual synchronization levels in humans

**DOI:** 10.1101/2022.11.24.517800

**Authors:** Marco Fuscà, Felix Siebenhühner, Sheng H. Wang, Vladislav Myrov, Gabriele Arnulfo, Lino Nobili, J. Matias Palva, Satu Palva

## Abstract

Moderate levels of synchronization of neuronal oscillations are essential for healthy brain dynamics. Synchronization levels exhibit large inter-individual variability the origins of which are unknown. Neuronal systems have been postulated to operate near a critical transition point or in an extended regime between disorder (subcritical) and order (supercritical phase) characterized by moderate synchronization and emergent power-law long-range temporal correlations (LRTCs). We investigated whether inter-individual variability in synchronization levels is explained by the individual position along the critical regime by analyzing magnetoencephalography (MEG) and intra-cerebral stereo-electroencephalography (SEEG) human resting-state data. Here we show that variability in synchronization levels exhibits a positive linear and quadratic relationships with LRTCs in healthy participants and brain areas. In the epileptogenic zone this correlation was negative. These results show that variability in synchronization levels is regulated by the individual position along an extended critical-like regime, with healthy brain areas tending to operate in its subcritical and epileptogenic areas in its supercritical side.

## Introduction

Brain activity is characterized by transient, long-range synchronized oscillations that play a fundamental role in regulating neuronal processing and communication across the brain (Fries, 2015; Hahn et al., 2019; Singer, 1999), which are essential to cognitive functions and behaviour (Fell & Axmacher, 2011; S. Palva & Palva, 2012, 2018; Schnitzler & Gross, 2005; Siegel et al., 2012; Thut et al., 2012). Healthy brain functioning can be achieved only at moderate levels of synchronization, while inadequate or excessive synchrony is characteristic to many brain disorders and associated with functional deficits (Ajramj et al., 2017; Bruining et al., 2020; Pusil et al., 2019; Uhlhaas et al., 2006; Hirvonen et al., 2018). Even among healthy subjects, however, there is a considerable variability in the mean levels of synchronization in large-scale brain networks (J. M. Palva et al., 2013; Smit et al., 2011; Wiesman et al., 2022). The factors underlying this variability and regulating synchronization levels are not well known.

The framework of brain criticality offers a putative explanation for this. The ‘critical brain’ hypothesis posits that neuronal systems operate at the critical point of a transition between disordered and ordered (subcritical and supercritical) phases, or between attenuating and amplifying activity propagation, respectively (Chialvo, 2010; Cocchi et al., 2017; Haldeman & Beggs, 2005; Levina et al., 2014; Plenz & Thiagarajan, 2007).

Operating at such a critical point gives rise to emergent power-law spatio-temporal correlations, intermediate mean levels of large-scale synchronization with large variance, and a range of functional benefits. At the experimental level, these can be indexed as power-law long-range temporal correlations (LRTCs) in fluctuations of local amplitudes (Linkenkaer-Hansen et al., 2001; J. M. Palva et al., 2013; Poil et al., 2012; Smit et al., 2011; Zhigalov et al., 2015) and power-law scaling of inter-areal ‘avalanche’ events (Beggs et al., 2007; Friedman et al., 2012; Haldeman & Beggs, 2005; Shew et al., 2011; Yang et al., 2012; Zhigalov et al., 2015), which together suggest that neuronal oscillations in vivo in humans and animal models indeed express critical-like dynamics. Theoretical work suggests that inter-areal synchronization, as a form of an emergent correlation, should also be dependent on criticality. Indirect support for this comes from results showing that the architecture of synchronization is indeed co-localized with that of LRTCs and avalanches (Zhigalov et al., 2017). Yet, despite the extensive work both on the synchronization and the power-law scaling of neuronal activity, there is little empirical evidence linking the actual strength of large-scale oscillatory synchronization with individual critical dynamics in the human brain.

LRTCs in oscillations amplitudes exhibit large variability across individuals, regions, and brain states implying that there may be a diversity of individual *operating points*, i.e., the positions of operation along the sub- to supercritical continuum around criticality (Linkenkaer-Hansen et al., 2001; Nikulin & Brismar, 2004; J. M. Palva et al., 2013; Smit et al., 2011; Zhigalov et al., 2015; 2017). Moreover, extending the original brain criticality hypothesis, recent theoretical studies suggest that human brains may not operate at a single critical point, but rather in an extended regime of critical-like dynamics known as the Griffiths phase (Moretti & Muñoz, 2013; Ódor & de Simoni, 2021). While there is yet little experimental support for this notion, it is in line with findings showing that different brain systems exhibit partially independent operating points (Zhigalov et al., 2017).

We hypothesized that individual variability in the operating point(s) of the functional brain systems in the critical regime would predict the individual variability in the synchronization levels and LRTCs. Brains have been predicted to operate very slightly subcritically to a critical point (Priesemann et al., 2014; Toker et al., 2022). We postulate here that rather than operating near a critical point, healthy brains operate on the subcritical side of an extended critical regime characterized with a diversity of individual operating points. Such an operating regime would yield the functional benefits of criticality and prevent the risks that excursions to the supercritical side would entail, i.e., runaway surges of excessive synchrony that characterize, e.g., epilepsy (Meisel et al., 2015; Monto et al., 2007).

If this were the case, the individual levels of inter-areal oscillatory synchronization would be positively correlated with LRTCs both across individuals and brain areas. We tested this hypothesis analyzing resting-state brain activity recordings of healthy subjects with non-invasive magnetoencephalography (MEG) and of subjects with drug-resistant epilepsy with intracranial stereo-electroencephalography (SEEG; Figure 1A). We then assessed the presence of LRTCs in the amplitude fluctuations using Detrended Fluctuation Analysis (DFA), which estimates the power-law scaling exponent of scale-free signals (Linkenkaer-Hansen et al., 2001). We calculated whole-brain pairwise inter-areal synchronization connectomes and derived cortical and individual measures of phase synchronization (*Figure 1B*). We then estimated whether synchronization at the whole-brain scale, and at the cortical region level, was correlated with the strength of LRCTs across individuals. We compared the results with modelling predictions to establish the position of individual brain dynamics in the critical regime.

**Figure 1.**
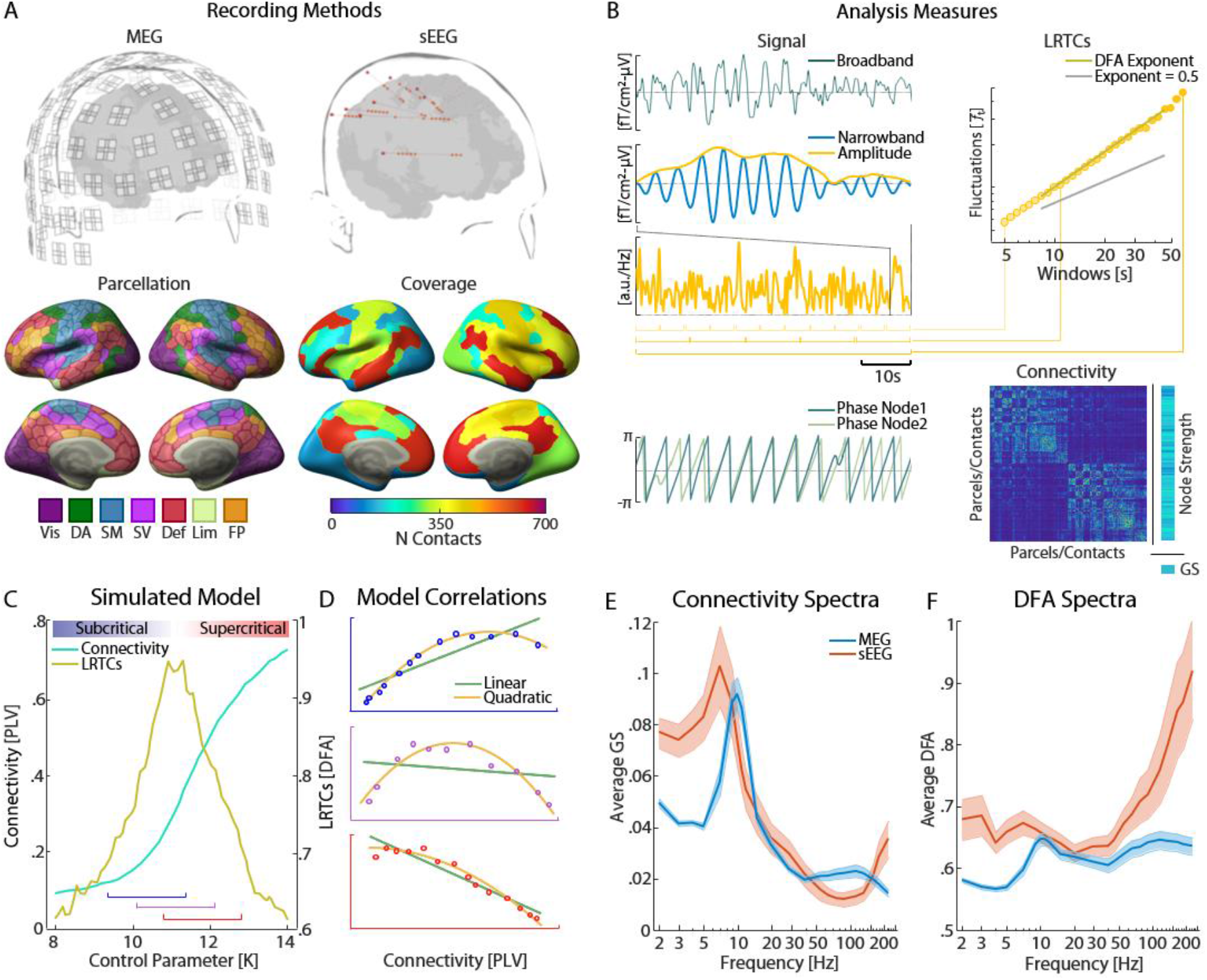
Study schematics. **A**. Schematic of recording with magnetoencephalography (MEG) and stereo-encephalography (SEEG). MEG sensor data (top left) was source-reconstructed to recover resting brain activity from 400 cortical parcels grouped according to functional connectivity (FC) systems (bottom left). Intracranial SEEG electrodes close to cortical grey matter from non-epileptic contacts were used and mapped to the same cortical parcellations as MEG (sample subject on top right, network contact coverage on bottom). **B**. MEG parcels and SEEG contact signals were filtered to obtain narrow-band amplitudes and phase time series (left) which were used to compute long-range temporal correlations (LRTCs) with Detrended Fluctuation Analysis (DFA; top right) and phase synchronization between brain areas. **C**. A computational Kuramoto model of critical synchronization dynamics visualizing the inter-dependence of synchronization and LRTCs measures shows that inter-areal synchronization increases monotonically with coupling (control parameter K), whereas LRTCs peak at the critical point. **D**. The relationship between synchronization and LRTCs increases linearly in the subcritical regime (top), quadratically around the critical point (middle), and decreases linearly in supercritical regime (bottom). **E**. Grand-average *GS* in MEG and SEEG data as a function of frequency. **F**. Same as in *E*. for DFA exponents. Shaded areas represent the 95% bootstrapped confidence intervals.

## Results

### Kuramoto model

We first used computational modelling to assess how the correlations of large-scale synchronization and LRCTs are determined by the operating point in the critical regime. We modelled local and large-scale neuronal oscillatory dynamics with a nested variant (Siebenhühner et al., 2020) of the Kuramoto model (Simola et al., 2022) with 100 regions containing 500 oscillators each. In this model coupling parameters *K* and *L* control the intra- and inter-regional, respectively, coupling strengths. The structural connectivity in local (nodal) networks is homogenous while the pairwise connectivity strengths between all cortical regions was based on white-matter axonal fiber counts estimated with structural diffusion tensor imaging (DTI). The model yielded synchronization dynamics both at local and large-scale network levels, which are directly comparable with the empirically observable time-series for each cortical parcel and their inter-areal interactions.

By increasing the within-node coupling, *K*, inter-areal synchronization estimated with the phase-locking value increased monotonically from low levels in the subcritical phase to near-perfect synchronization in the supercritical phase, while the DFA scaling exponents peaked around the phase transition, i.e., in the critical regime, and fell off sharply to the left and right (*Figure 1C*). Therefore, DFA exponents were correlated positively with synchronization in the subcritical slope of the critical regime and negatively in the supercritical slope (Figure 1D). The control parameter *K* here is conceptually comparable with excitation-inhibition ratio (E/I), or “neuronal gain”, which in other models comprise the functional net effects of inhibitory and excitatory synaptic, cellular, biophysical, and microcircuit mechanisms at the nodal level (Deco & Kringelbach, 2017).

Prior research has repeatedly shown that both large-scale synchronization and local LRTCs exhibit considerable systematic inter-individual variability (Nikulin & Brismar, 2005; J. M. Palva et al., 2013) that is trait-like and partly heritable (Leppäaho et al., 2019; Linkenkaer-Hansen et al., 2007). We posit here that this variability is driven by inter-individual differences in the operating point and positioning in the critical regime, *i.e*., in the physiological control parameters regulating brain criticality *in vivo*. The brains are thought to operate in the subcritical side of the critical point (Priesemann et al., 2014). This would give rise to inter-individual variability where both synchronization and LRCTs monotonically increase and have mainly a positive linear correlation (*Figure 1D*, top panel). If the population would operate around the critical point, correlation between synchronization and LRCTs should exhibit a quadratic correlation with an inverted-U shape (*Figure 1D*, middle panel), indicated by a negative quadratic coefficient. In supercritical dynamics, possibly a characteristic of an epileptic state (Meisel, 2016; Meisel et al., 2015; Monto et al., 2007), the correlations between synchronization and LRTCs should again have a linear relationship, but with a negative slope (*Figure 1D*, bottom panel).

### Group averages of synchronization and LRTCs

To assess oscillatory brain dynamics at the whole-brain level, we computed pairwise phase synchronization for all narrow-band frequencies between 400 cortical parcels (Schaefer et al., 2018) from source-reconstructed MEG data (192 sessions from 52 healthy participants) and between all non-epileptogenic contact-pairs of SEEG data (57 epileptic subjects, 1 session each). To estimate phase synchronization, we used the Phase Locking Value (PLV) for SEEG and the weighted Phase Lag Index (wPLI) (Vinck et al., 2011) for MEG source-data because PLV in MEG is inflated by artefactual zero-phase lagged synchronization (see *Materials and Methods*). Graph strength (*GS*) (Bullmore & Sporns, 2009), defined as the mean connectivity, was used to assess the individual level of global synchronization.

In MEG (*N* = 192 sessions from 52 subjects), the grand-averaged *GS* peaked at the alpha frequency band (8–14 Hz), while in SEEG (*N* = 57 patients, 1 session each), *GS* peaked slightly lower at theta-alpha (5–10 Hz; *Figure 1E*) in line with previous studies showing a shift from alpha to theta in SEEG (Arnulfo et al., 2020; Siebenhühner et al., 2020; Wang, 2021). We next assessed the presence of LRTCs in the narrow-band oscillation amplitude envelopes by estimating the scaling exponent with DFA (Hardstone et al., 2012). We obtained individual mean values also for DFA exponents (mean DFA) by averaging across parcels for MEG and contacts for SEEG data for each session. In MEG, the group-averaged mean DFA showed a well-delineated alpha band peak as seen in *GS* above and a broader gamma band peak (50–100 Hz). In SEEG, group-averaged mean DFA peaked in the theta-alpha band (5–10 Hz) as well as in delta band (2–4 Hz) and exhibited a monotonic near-linear increase in gamma frequencies (*Figure 1F*).

### Synchronization and LRTC exponents are positively correlated across individuals

To study whether interindividual variability could be explained by the brain criticality hypothesis, we investigated whether *GS* and DFA values would be correlated and co-vary across subjects. In MEG, the correlations between *GS* and mean DFA were significant in all frequencies above 4 Hz (Pearson correlation test, FDR-corrected, *p_FDR_* < 0.01 for 4–5 Hz and *p_FDR_* < 10^−7^ for the higher frequencies, reaching *p_FDR_* < 10^−25^ at 7 Hz; *Figure 2A*). In SEEG, the correlations between *GS* and mean DFA were weaker than in MEG in higher frequencies, but significant in theta-alpha (Pearson correlation test, 4–13 Hz, *p_FDR_* < 0.01 for 4 and 10 Hz) and “ripple” high gamma (>165 Hz) bands (Pearson correlation test, *p_perm_* < 0.05, *Figure 2B)*. We then also computed correlations between *GS* and DFA values also within the canonical frequency bands that had been confirmed using Louvain clustering (*Supplementary Figure 1)* and observed significant linear positive correlations in all frequencies in MEG and in theta (θ, 4–7 Hz), alpha (8–12 Hz) and beta (β, 15–29 Hz) in SEEG (*Supplementary Figure 2*). We further confirmed that results did not depend on metrics using PLV *GS* for MEG and wPLI for SEEG (*Supplementary Figure 3*). As predicted by our modeling results, the positive correlations between *GS* and mean DFA indicate that brain dynamics operate on the subcritical side of a critical regime.

**Figure 2.**
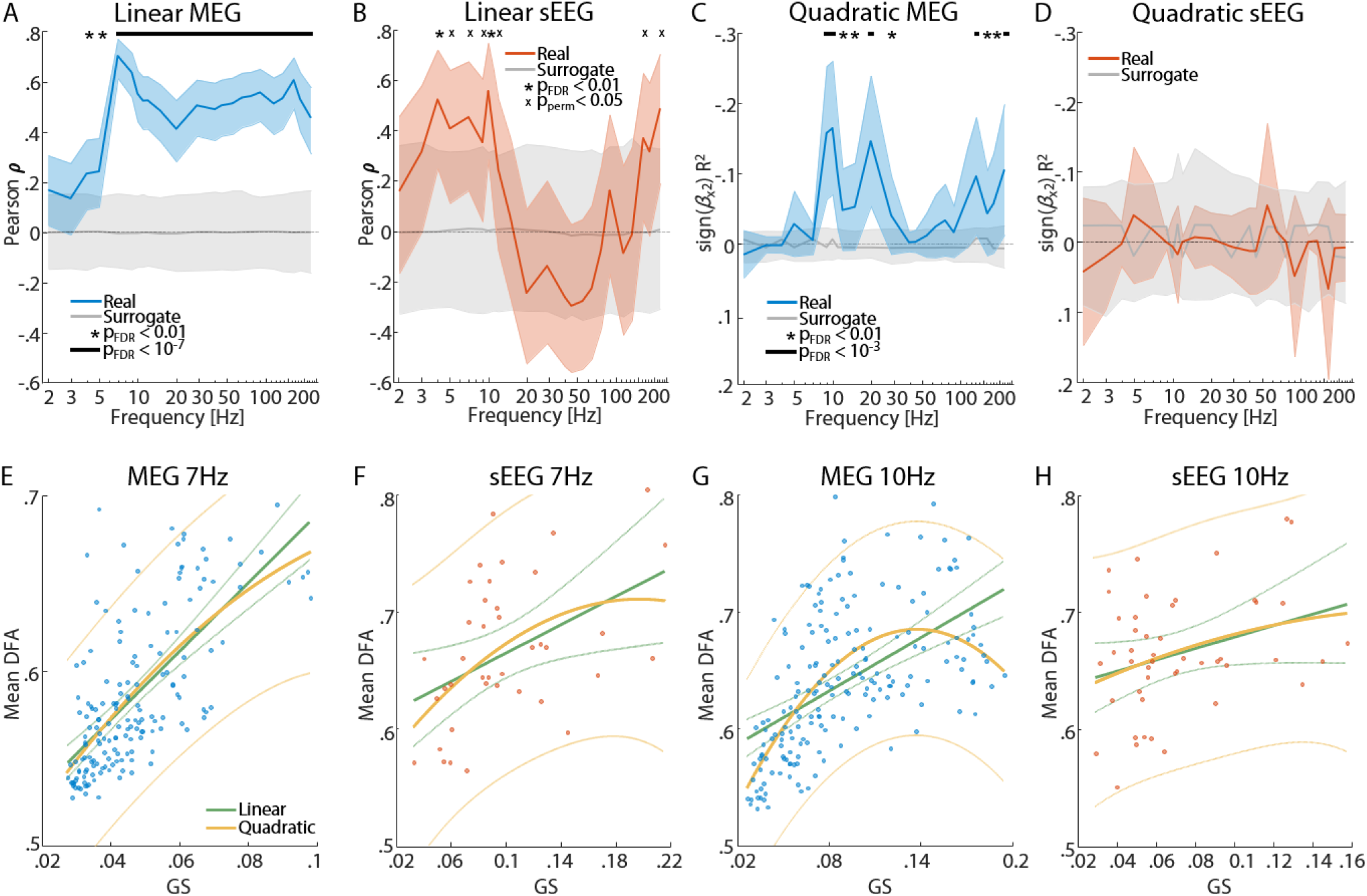
Correlations between global synchronization GS and LRTCs. **A**. Correlation of individual set’s *GS* and mean DFA values (Pearson correlation test) for MEG data. Shading shows the 95% confidence intervals and gray areas the 2.5-97.5^th^ percentiles of the surrogate coefficients distribution. The asterisks indicate *p_FDR_* < 0.01 after FDR correction; the black line at the top indicates *p_FDR_* < 10^−7^. **B**. Same as in *A*. but for SEEG. Significant p-values that do not survive FDR but are above the surrogate 97.5^th^ percentile are marked by *X*. **C**. Partial quadratic correlations of *GS* and mean DFA (See Methods) for MEG data as in *A*; black line at the top is *p_FDR_* < 10^−3^; asterisks are *p_FDR_* < 0.01. **D**. Same as in *C* for SEEG. **E**. Scatterplot of *GS* and mean DFA values at 7 Hz with each dot representing a session. Solid lines represent linear (green) and quadratic (yellow) fits, with the faint dotted side lines showing the 95% prediction bounds. **F**. Same as in *E*. for SEEG. **G.H**. Same as in *E*. and *F*. for 10 Hz.

Our modeling results further implied that a system situated in the vicinity of the critical transition would be evidenced by quadratic correlations in addition to linear ones (see *Figure 1D*). We thus estimated also quadratic trends and their direction with partialed-out linear influences between *GS* and mean DFA, using the R^2^ regression statistic multiplied with the sign of the quadratic coefficient. At the critical point, the quadratic coefficient should be negative, denoting a peak, *i.e*., a concave curve with an inverted-U shape. Significant correlations with a negative quadratic coefficient between *GS* and mean DFA were indeed observed in MEG data in alpha (8–12 Hz, peak frequency at 10 Hz with *p_FDR_* < 10^−6^), beta (17–30 Hz) and high-gamma (>135 Hz; *Figure 2C*) bands. In SEEG data, on the other hand, there were no significant quadratic correlations between *GS* and mean DFA in any of the frequencies (*Figure 2D*). We then plotted the correlation of *GS* and mean DFA values across subjects for the frequencies with the highest correlations, observing predominantly linear relationships with an additional quadratic component in MEG (*Figure 2E-H*). Quadratic correlations between *GS* and DFA values were not found in the canonical frequency bands (*Supplementary Figure 2*), but were reproduced in MEG using PLV instead of wPLI (*Supplementary Figure 3*).

As both synchronization and LRTCs have been shown to be trait-like phenomena, we further investigated whether their correlations would exhibit high retest reliability as required by a trait-like phenomenon. Using the Gauge Repeatability method (Burdick et al., 2005) we confirmed that both individual *GS* and DFA values (*Supplementary Figure 4A-C*) and their correlations (*Supplementary Figure 4D-E)* had significant retest reliability and capacity.

### Synchronization and LRTC exponents are correlated across brain regions

To get insight into the anatomy of these correlations, we estimated the correlation of synchronization and DFA exponents across subjects separately for each parcel by estimating a mean nodal synchronization using Node Strength (*NS*; with MEG parcels or SEEG contacts being the nodes) and its correlations with the DFA exponents of the node within the canonical frequency bands. In order to be able to compare results between MEG and SEEG, SEEG data were collapsed into the same atlas of 400 parcels (Schaefer et al., 2018) that was used for MEG. Positive linear correlations between NS and DFA exponents were significant in most frequencies but strongest in the alpha and gamma bands within all functional subsystems in MEG data (*Figure 3A, Supplementary Figure 5A*). In SEEG, significant positive linear correlations were observed in the delta and alpha frequency bands, while negative correlations were present in low and high gamma bands, but again positive in ripple-high gamma frequencies (*Figure 3B, Supplementary Figure 5B*). In MEG, linear correlations were widespread throughout the cortex, highest in dorsal areas (*Figure 3C*). In SEEG, positive linear correlations in theta and alpha bands were strongest and lateral temporal and parietal regions belonging to the Default Mode (DM) network where most contacts were located, and also found in the prefrontal cortex (PFC, *Figure 3D*).

**Figure 3.**
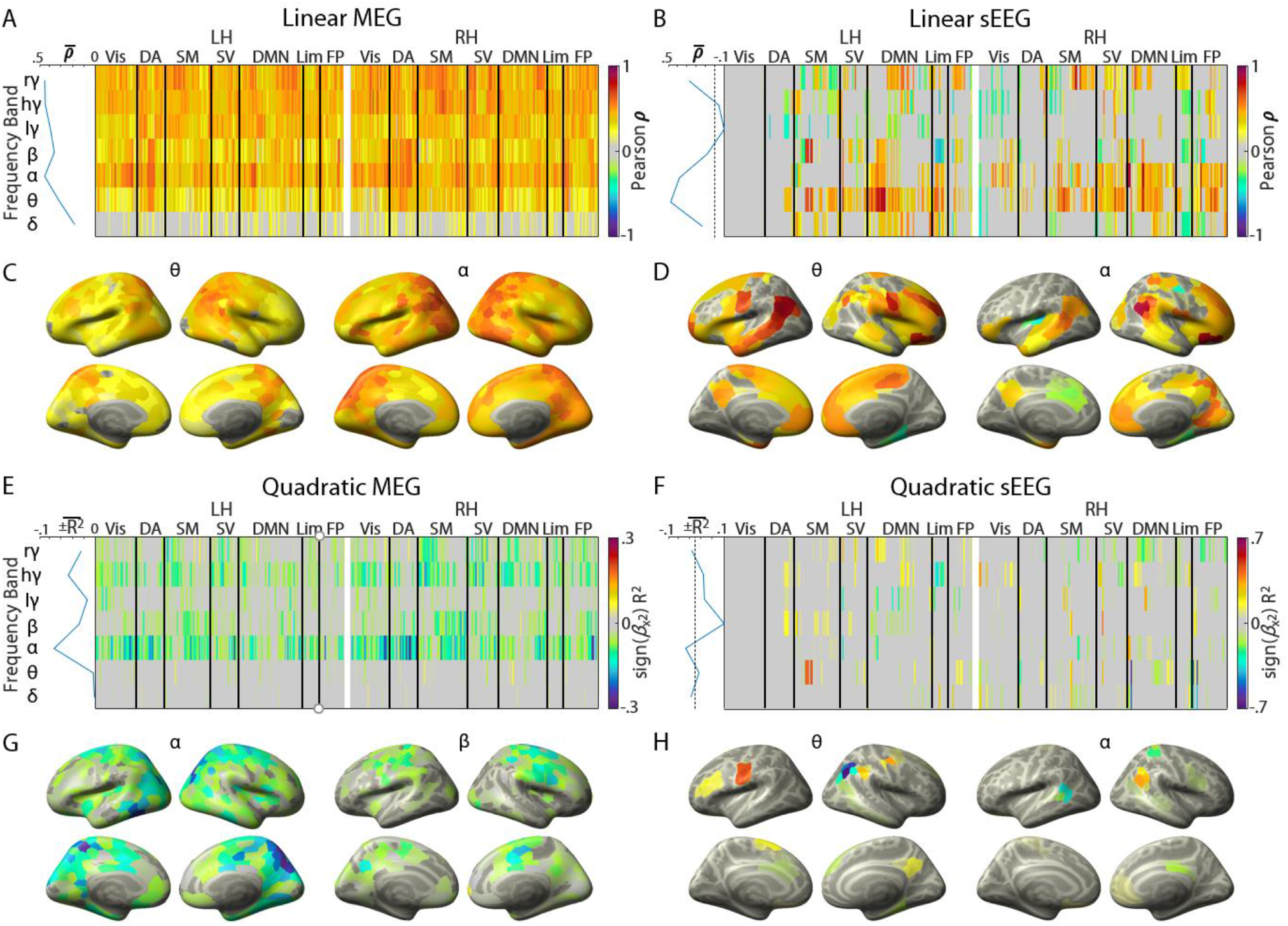
Correlations of local synchronization *NS* and LRTC exponents. **A**. Correlation of *NS* and mean DFA values estimated for each parcel (Pearson correlation test) in MEG data for left and right hemisphere (LH and RH) averaged over canonical frequency bands. The blue line on the left shows the mean correlation across significant parcels for each band. **B**. Same as in *A*. for SEEG. **C**. Cortical topographies of the correlation between MEG *NS* and DFA for theta and alpha frequency bands. **D**. Same as in *C*. for SEEG. **E-H**. Same as above for partial quadratic correlation. Frequency bands: δ 2-4 Hz; θ 4-8 Hz; α 9-12 Hz; β 15-29 Hz; lγ 40-65 Hz; hγ 77-135 Hz; rγ 165-225 Hz. Parcels networks: *Vis*: Visual; *DA*: Dorsal Attention; *SM*: Somatomotor; *SV*: Saliency/Ventral Attention; *DMN*: Default Mode Network; *Lim*: Limbic; *FP*: Frontoparietal/Control.

Negative quadratic correlations (again obtained after regressing out the linear trend) were observed from alpha to gamma bands in MEG (*Figure 3E, Supplementary Figure 5C*) and with lower power in the theta and alpha band in SEEG data (*Figure 3F, Supplementary Figure 5D*). Negative quadratic coefficients in MEG indicated that the quadratic trends were again all concave (opening down, peaking with a parabolic maximum) and hence indicated brain dynamics being located close to the critical point.

In SEEG, quadratic correlations were characterized by both negative and positive coefficients, the latter especially in beta and gamma frequency bands. Negative quadratic correlations in MEG were widespread in frequencies above alpha, strongest in posterior regions in alpha and in somatomotor network in the beta band (*Figure 3G*). In SEEG, significant correlations were sparser, with a strong negative quadratic correlation in DMN parietal parcels in the theta band (*Figure 3H*, other bands in *Supplementary Figure 1*). These results were reproduced also using individual wavelet frequencies (*Supplementary Figure 6*) and had significant retest reliability *(Supplementary Figure 4F-I)*.

### Negative correlations in contacts within the Epileptogenic Zone (EZ)

The results thus far showed that healthy brain dynamics are predominantly characterized by linear positive correlation between synchrony and DFA with a quadratic trend, which is in line with the notion of human brains operating on the subcritical side of an extended critical regime. Epilepsy has been associated with excessive excitation, hyper-synchrony (Arnulfo, Hirvonen, et al., 2015; Parish et al., 2004), and altered DFA exponents (Auno et al., 2021; Meisel et al., 2015; Poil et al., 2012). Hence, we predicted that contacts in the EZ (*Figure 4A*) should be associated with supercritical dynamics, in contrast to healthy brain activity operating in the subcritical regime, comparatively visible in the non-epileptogenic (non-EZ) contacts in the previous analyses.

**Figure 4.**
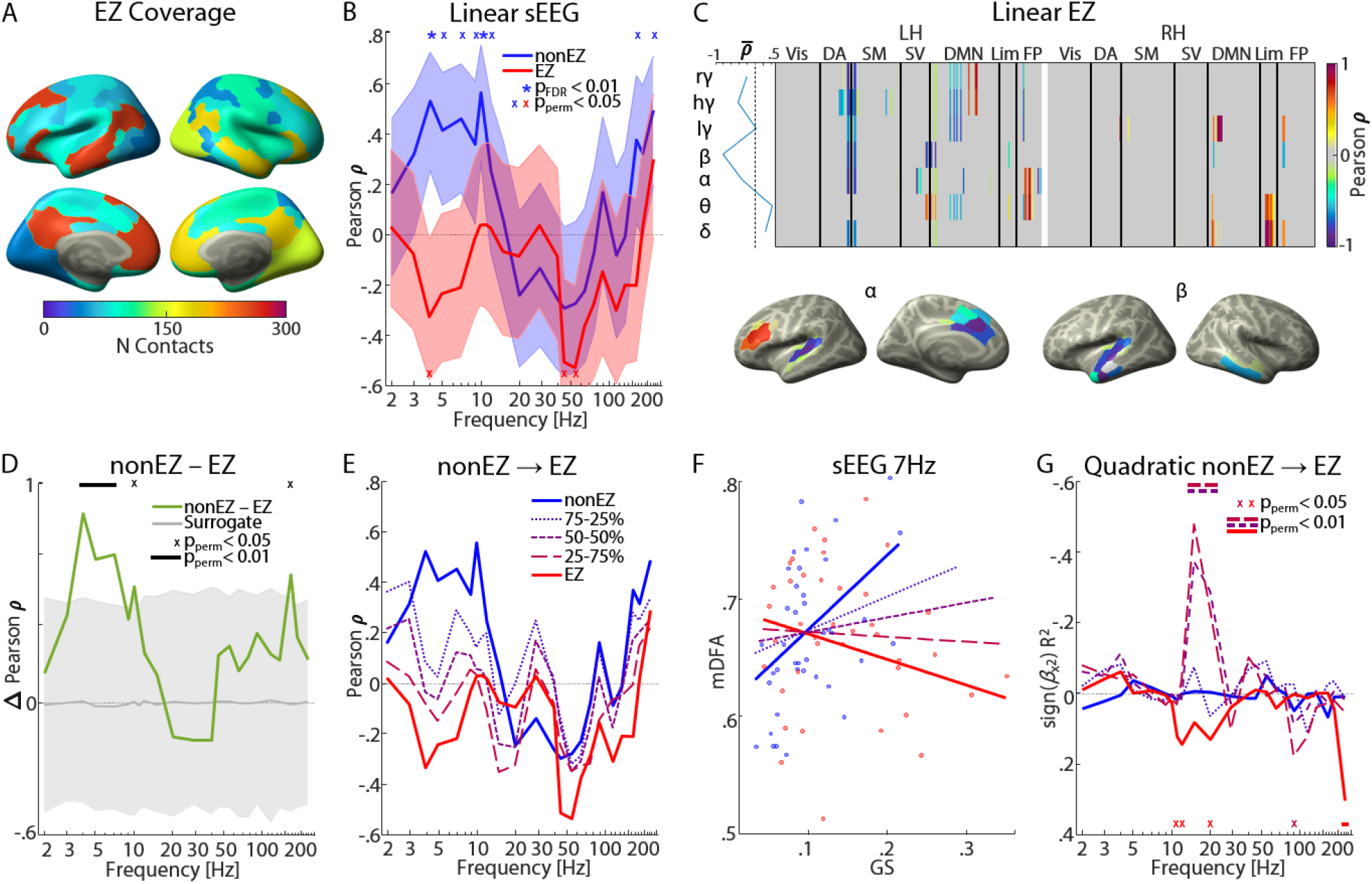
Correlations of synchronization and LRTCs for epileptic zone (EZ) SEEG contacts. **A**. Cortical distribution of EZ contacts across the functional system parcellation of the Schaefer atlas. **B**. Linear correlation between *GS* and mean DFA, with non-EZ correlations in blue (same as in *Figure 2B*) and EZ correlations in red, with shaded errorbar areas representing 95% confidence intervals. Color-coded asterisks at the top indicate *p_FDR_* < 0.01 and blue and red Xs at the top and bottom indicate *p_perm_* < 0.05. **C**. Linear correlation of *NS* and mean DFA for OZ contacts a. **D**. Difference correlation coefficients between non-EZ and EZ contacts tested against a surrogate difference distribution obtained with case-resampling permutations. **E**. Same as in *B*, but with intermediate steps gradually increasing the percentage of EZ contacts and decreasing that of non-EZ ones, showing the progressive emergence of pathological dynamics. **F**. Scatterplot of *GS* and mean DFA values at 7 Hz each dot representing data for one subject. The dashed lines represent the intermediate steps same as in *B*. **G**. Same as in *E*. for partial quadratic correlations.

We thus investigated correlations between synchrony and LRTCs for EZ contacts in SEEG. As hypothesized, the correlations between subjects’ *GS* and mean DFA were negative and significant at 4 Hz and 45–55 Hz for EZ contacts (*p_perm_* < 0.05, *Figure 4B*). Similarly, negative coefficients were also seen at the parcel-level correlations of NS and DFA exponents (*Figure 4C*), with significant correlations in alpha in left temporal and cingulate areas of the DMN, with the beta band also showing negative correlations in the right temporal cortex, in which many EZ electrodes were located. The difference in mean correlation values between non-EZ and EZ was significant between 3 and 10 Hz (peaking at 4 Hz with *p_perm_* < 10^−7^, *Figure 4D*) and at 165 Hz (*p_perm_* < 0.05, *Figure 4D*).

We then repeated the group-level correlations in three intermediate steps in which we increased the number of EZ contacts and decreased the number of non-EZ contacts to show the gradual change from positive to negative linear coefficients as an index of the emergence of pathological dynamics. We observed a gradual shift from positive to negative linear correlations in the theta and alpha bands (*Figure 4E*). This trajectory was also observed in the single frequency scatterplots of *GS* and mean DFA with linear fits gradually crossing over to the negatives (*Figure 4F*). Partial quadratic correlations (with the linear component regressed out) displayed significant negative quadratic correlations in the intermediate steps, peaking at the beta frequency band (with *p_perm_* < 10^−4^ for 15 Hz of the 25-75% condition), and positive quadratic correlations in EZ-only contacts in single frequencies in the alpha (11–12 Hz), beta (20 Hz) and ripple-high gamma bands (225 Hz, all at *p_perm_* < 0.05, *Figure 4G*).

## Discussion

The emergence of power-law inter-areal and temporal correlations (Chialvo, 2010) in brain activity has been proposed to be attributable to brain criticality and thus regulated by its underlying physiological control parameters. Both synchronization and LRTCs in oscillatory amplitude emerge at the critical point both in modelling (Botcharova et al., 2014; Gray & Robinson, 2007; Larremore et al., 2011; Levina et al., 2014; Martinello et al., 2017; Moretti & Muñoz, 2013, in vivo (Fontenele et al., 2019), and in vitro (Heiney et al., 2021) studies.

However, there is a considerable variability in the mean strength of synchronization networks and in the mean LRTCs (J. M. Palva et al., 2013; Simola et al., 2017; Smit et al., 2011; Wiesman et al., 2022; Zhigalov et al., 2017). Yet, neither the relationship between synchronization and LRTCs, nor the reasons causing the large inter-individual variability, are well understood. As oscillatory synchronization is a fundamental property of brain networks, critical for information processing and cognition, and relevant for multiple brain disorders (Fries, 2015; Hahn et al., 2019; S. Palva & Palva, 2012, 2018; Siegel et al., 2012; Thut et al., 2012), understanding the underpinnings of individual levels of synchronization is of high importance.

Here, we tested whether large inter-individual variability in both synchronization dynamics and oscillation LRTCs would be explained by the individual’s position in the critical regime, *i.e*., by the individual operating point. We took advantage of the large inter-individual variability and the differential dependence of these measures from the critical point. We show here that inter-areal synchronization is positively correlated with the scaling exponents of LRTCs in neuronal oscillations together with a small but significant quadratic trend. The positive correlation suggests that most subjects operate in the subcritical side of an extended critical regime, while the observation of a significant quadratic relationship suggests that some subjects operate around the peak of this regime, *i.e*., very near the critical tipping point.

Our results demonstrate that individual variability in synchronization levels and in LRCTs are linked through a relationship predicted by the framework of brain criticality. These findings are in line with the framework where critical-like dynamics emerge across a “stretched” or extended critical regime, also known as the Griffiths phase (Moretti & Muñoz, 2013; Ódor & de Simoni, 2021), as opposed to criticality being constrained to a singular point in the space of control parameter. To the best of our knowledge, these findings thus constitute the first empirical evidence towards the discovery of Griffith’s-phase-like dynamics in the brains *in vivo*. Moreover, these results suggest that human brains do not exactly operate in a ‘subcritical’ regime (Priesemann et al., 2014; Wilting & Priesemann, 2019) near the critical point, but rather within an extended critical regime even if on its subcritical side. This distinction is fundamental because only the critical regime enables the emergence of power-law spatiotemporal structures and the numerous functional benefits that the brains may gain by operating with critical-like dynamics.

As both connectivity and LRTCs are test-retest reliable (Candelaria-Cook et al., 2022; Nikulin & Brismar, 2004; Wiesman et al., 2022), heritable (Leppäaho et al., 2019; Linkenkaer-Hansen et al., 2007; Nikulin & Brismar, 2005), and influenced by genetic polymorphisms (Simola et al., 2022), we postulate that the individual’s position in the critical regime is a phenotypic trait that predicts individuals’ brain dynamics and behaviors. We postulate that individual variability in critical dynamics could underlie variability in cognitive abilities and personality traits (J. M. Palva et al., 2013) given that operating at critical dynamics is thought to maximize the dynamic range of the system, sensitivity to external and internal perturbations, transmission, storage, and computational capacity (Beggs et al., 2007; Chialvo, 2010; Larremore et al., 2011; S. Palva & Palva, 2018; Shew et al., 2011; Toker et al., 2022).

In contrast to healthy brain activity, epilepsy has been associated with excessive excitation leading to aberrant pathological brain dynamics (Arnulfo, Hirvonen, et al., 2015; Bartolomei et al., 2013; Parish et al., 2004) and episodes of abnormal hyper-synchronous activity (Monto et al., 2007). Since brain criticality is primarily thought to be controlled by the finely balanced E/I ratio, where excessive excitation leads to super-critical dynamics (Plenz & Thiagarajan, 2007; Poil et al., 2012; Shew et al., 2009; Toker et al., 2022), we investigated whether epilepsy would be characterized by dynamics in the supercritical regime. This would be marked by negative correlations between synchronization and DFA exponents (see Fig. 1). We found that the EZ contacts were indeed associated with a shift towards negative correlations between synchronization and LRTCs. This constitutes new empirical evidence for epilepsy being associated with supercritical brain dynamics (Meisel, 2016; Meisel et al., 2015) that may be the underlying cause for hyper-synchronous activity and predisposition of these brain networks to generate epileptic seizures.

In SEEG, in the nominally healthy “non-EZ” contacts outside of the epileptogenic zone, there were no quadratic correlations indicative of dynamics near the critical tipping point. This could be caused by a push-pull mechanism where the brain compensates for increased excitability and super-critical dynamics via an excess of inhibition, which then drives the healthy brain regions in epileptics towards sub-criticality, or by antiepileptic drugs. Synchronization and LRTCs are aberrant also in multiple brain other diseases (Ajramj et al., 2017; Auno et al., 2021; Bajo et al., 2012; Bartolomei et al., 2013; Bruining et al., 2020; Linkenkaer-Hansen et al., 2005; Meisel et al., 2015; Monto et al., 2007; Parish et al., 2004; Pusil et al., 2019; Smit et al., 2011; Uhlhaas et al., 2006). Taken that brain criticality is primarily thought to be controlled by the E/I ratio, where an imbalance of E/I or connectivity leads to sub- or super-critical dynamics being modulated by polymorphism in neuromodulatory genes (Plenz & Thiagarajan, 2007; Poil et al., 2012; Shew et al., 2009, 2011; Simola et al., 2022) and that pathological human brain activity is associated with changes in brain E/I balance (Bajo et al., 2012; Pusil et al., 2019; Uhlhaas et al., 2006), pathological synchronization dynamics (hypo- or hyperconnectivity) could emerge via modulations of brain critical dynamics.

## Conclusions

Our results demonstrate that variability in synchronization levels is regulated by the individual position along an extended critical regime so that healthy brain areas operate in its subcritical and epileptogenic areas in the supercritical side.

## Materials & Methods

### Modeling

A nested Kuramoto model (Siebenhühner et al., 2020) was used to simulate coupled neuronal populations dynamics with LRTCs and to investigate observable correlations between synchronization and DFA exponents. We adapted a two-layer nested model consisting of 100 regions/nodes, each containing a conventional Kuramoto population of oscillators (N = 500). The oscillators’ phases were averaged to derive a nodes’ time series, whose absolute values were then taken to obtain the Kuramoto order parameter, corresponding to their amplitude. Pairwise connection weights were proportional to structural connectivity values obtained by MRI diffusion-tensor imaging and averaged to the Schaefer atlas with 100 parcels (Schaefer et al., 2018). Thus, we were able to model both local interactions in smaller regions (corresponding to cortical MEG parcels or SEEG contacts), and their large-scale synchronization (corresponding to inter-areal connectivity across the whole brain). This model contained two control parameters, K as the strength of internal phase coupling within regions, and L as the strength of inter-regional coupling. We estimated LRTCs of regions’ time series using DFA, and synchronization between regions using the PLV (see below).

### Acquisition of MEG and MRI Data

We recorded MEG data from 52 healthy participants (age: 31 ± 9.2, 27 male) during a 10-minute eyes-open resting-state session with a Vectorview/Triux (Elekta-Neuromag/MEGIN, Helsinki, Finland) 306-channel system (204 planar gradiometers and 102 magnetometers) at the Bio-Mag Laboratory, HUS Medical Imaging Center, Helsinki. Overall, 192 sessions of MEG data were obtained, with participants contributing on average 3.7 ± 4 sessions each. Participants were instructed to focus on a cross on the center of the screen in front of them. Bipolar horizontal and vertical electrooculography (EOG) were recorded for the detection of ocular artifacts. MEG and EOG were recorded at 1 kHz sampling rate. For each participant, T1-weighted anatomical MRI scans (MP-RAGE) at a resolution of 1 × 1 × 1 mm with a 1.5-Tesla MRI scanner (Siemens, Munich, Germany) were obtained at Helsinki University Central Hospital for head models and cortical surface reconstruction. The study protocol for MEG and MRI data was approved by the Coordinating Ethical Committee of Helsinki University Central Hospital (ID 290/13/03/2013), written informed consent was obtained from each participant prior to the experiment, and all research was carried out according to the Declaration of Helsinki.

### Cortical parcellation and source model

Volumetric segmentation of MRI data, flattening, cortical parcellation, and neuroanatomical labeling with the 400-parcel Schaefer atlas (Schaefer et al., 2018) was carried out with the FreeSurfer software (Fischl, 2012). The MNE software (Gramfort et al., 2014) was then used to create cortically constrained source models, for MEG–MRI colocalization, and for the preparation of the forward and inverse operators. The source models had dipole orientations fixed to pial-surface normals and a 5-mm interdipole separation throughout the cortex, which yielded around 5000-8000 source vertices per hemisphere.

### MEG data preprocessing and filtering

Temporal signal space separation (tSSS) (Taula & Simola J, 2006) in the Maxfilter software (Elekta-Neuromag) was used to suppress extracranial noise from MEG sensors and to interpolate bad channels. We used independent components analysis adapted from the Fieldtrip toolbox (Oostenveld et al., 2011) to extract and identify components that were correlated with ocular artifacts (identified using the EOG signal), heartbeat artifacts (identified using the magnetometer signal as a reference), or muscle artifacts.

### MEG source localization

We computed noise covariance matrices (NCMs) using the preprocessed MEG data time series filtered with finite-impulse-response (FIR) filters at 151–249 Hz, averaged across 10 s time-windows. NCMs were used for creating one inverse operator per session with the MNE software and the dSPM method with regularization parameter λ = 0.11 (Gramfort et al., 2014). We then estimated “vertex fidelity” to obtain fidelity-weighted inverse operators that reduce the effects of spurious connections resulting from source leakage, and collapsed the inverse-transformed source time series into parcel time series in a manner that maximizes the source-reconstruction accuracy as in (Korhonen et al., 2014; Rouhinen et al., 2020; Siebenhühner et al., 2020). For each parcel pair (edge) we also computed “cross-parcel phase-locking” of the reconstructed simulated time-series, reflecting cross-parcel signal mixing, and excluded parcels and edges with low fidelity and high cross-parcel phase-locking, using individual thresholds to retain for each subject the top 90% parcels by fidelity and the bottom 95% of edges by cross-parcel mixing (14.9 ± 0.2% of parcels and 14.1 ± 0.1% of edges rejected on average per set).

### Acquisition of SEEG Data

We recorded stereo-EEG neuronal signals from 68 drug-resistant focal epileptic patients (age: 30 ± 9.4, 38 male) during the clinical assessment of the epileptogenic focus before its surgical ablation at the “Claudio Munari” Epilepsy Surgery Centre in the Niguarda Ca’ Granda Hospital, Milan. Intracranial monopolar (with contacts sharing the reference to a single white-matter contact) local-field potentials were acquired from brain tissue with platinum–iridium multi-lead electrodes. Between 8 to 15 contacts, each 2 mm long, 0.8 mm thick and with an inter-contact border-to-border distance of 1.5 mm (DIXI medical, Besancon, France), were present in each penetrating shaft, with the amounts of electrodes and their anatomical positions varying according to surgical requirements (Cardinale et al., 2013). Each subject had 17 ± 3 (range 9-23) shafts with a total of 153 ± 20 electrode contacts on average. The electrode positions were localized after implantation using CT scans and the SEEGA automatic contact localization. Structural MRIs were recorded before implantation and colocalized with postimplant CT scans using rigid-body coregistration (Arnulfo, Narizzano, et al., 2015). Individual patients’ contacts were assigned to parcels of the Schaefer atlas (Schaefer et al., 2018). We acquired an average of 10 min of uninterrupted spontaneous resting-state activity with eyes closed with a 192-channel SEEG amplifier system (Nihon-Kohden Neurofax EEG-1100) at a sampling rate of 1 kHz. Patients gave written informed consent for participation in research studies and for publication of results pertaining to their data. The ethical committee of the Niguarda Hospital, Milan, approved this study (ID 939) which was performed according to the Declaration of Helsinki.

### Filtering and preprocessing of SEEG data

Defective contacts that demonstrated non-physiological activity (1.3 ± 1.2, range 0–10) were excluded from analyses, and 3 subjects in which more than 50% of contacts were defective were discarded from further analyses. We referenced SEEG electrodes in grey matter to the closest contacts in white matter (Arnulfo et al., 2014) which yields signals with more accurate phase estimates. Time series were FIR-filtered with a cutoff at 440 Hz and notch FIR filter was used for removing 50 Hz line noise and its harmonics. Temporal windows of 500 ms containing mean activity above 5+ SD in > 10% of cortical contacts and at least 25% of the narrow-band frequencies were rejected as potentially epileptogenic.

In clinical literature, epileptogenic zone is defined as the “minimum amount of cortex that need to be removed to produce seizure freedom”, while the seizure propagation network is defined as “the set of regions that participate in the propagation of epileptic activity during seizure onset”.

In this work, the epileptogenic zone and seizure propagation networks were identified by clinical expert analyses of the clinical data and quantitative diagnostic imaging data including SEEG traces, which were confirmed later by another expert (Cossu et al., 2015).

Most of the analyses presented in this work included only contacts from non-epileptic cortical regions, except for the Epileptic Contacts Correlations below, for which cleaning was performed as described above. Rejection of ictal activity, interictal events, and artefacts was also performed as before in order to avoid introducing obviously hypersynchronized dynamics (Arnulfo et al., 2020) and biasing the criticality assessment (Wang et al., 2022). Subjects who had more than half of their contacts in these areas (8 subjects) were additionally excluded from the normal correlations, as were contacts with more than 5% of segments containing epileptic events. For healthy contact analyses, we thus included data from 57 subjects, with a total of 4453 non-EZ contacts (average per subject 78 ± 19, range 41-124).

For the Epileptic Contacts Correlations, we used the contacts defined as being epileptogenic – i.e., contacts that were located within the EZ or were part of the seizure propagation network. Subjects discarded for having too many EZ contacts were included in this analysis, whereas subjects who had < 11 epileptogenic contacts were excluded. Thus, both analyses had 57 subjects, with 49 common to both non-EZ and EZ analyses, 8 only included in non-EZ, and 8 only included in EZ analysis. The total number of EZ contacts was 1725 (average per subject 30 ± 17.5, range 11-79).

### Analysis of inter-areal synchronization

Both cleaned MEG and SEEG data were filtered into complex-valued narrowband time series using Morlet wavelets (*m* = 5 width) with logarithmically increasing center frequencies ranging from 2 to 225 Hz. We estimated inter-areal functional interactions for each contact and parcel pair, but excluding the edges of adjacent contact pairs with distance < 2 cm, for each frequency and for each session, using both the Phase-Locking Value (PLV) (Lachaux et al., 1999) and the weighted Phase Lag Index (wPLI) (Vinck et al., 2011) which, unlike PLV, is insensitive to zero-phase lagged synchronization caused by source-mixing in MEG.

### Detrended Fluctuation Analysis

We used Detrended Fluctuation Analysis (DFA) (Linkenkaer-Hansen et al., 2001) to estimate monofractal scaling exponents of neuronal LRTCs that typically vary between 0.5 and 1 (Linkenkaer-Hansen et al., 2001, 2005; Meisel et al., 2015; Monto et al., 2007; Nikulin & Brismar, 2004, 2005). DFA was carried out in the Fourier domain (Nolte et al., 2019) with a Gaussian weight function used for detrending and using 25 log-linear windows from 5 s – 56 s. The fluctuations were fitted with a robust linear regression with a bisquare weight function to obtain the DFA exponents, all with negligible fit error. DFA exponents were computed for all contacts of all SEEG subjects and all parcels of all MEG sets, and for each narrow-band frequency.

### Average connectivity and criticality metrics

To assess the general level of synchrony of each node (contacts in SEEG and reconstructed sources in MEG), we used graph theory (Bullmore & Sporns, 2009). Node Strength (*NS*) of pairwise connectomes was obtained for each node by averaging the strength of edges, that is the mean synchronization of that node with all the other nodes. For each participant, all the cortical *NS* values were then averaged again to estimate the Graph Strength (*GS*) for each frequency and DFA scaling exponents of all nodes to acquire a mean DFA. We then removed outliers > 3 SD from the median. As inter-day variability was non-neglectable for violations of independence (*Supplementary Figure 4*; also see Candelaria-Cook et al., 2022; Wiesman et al., 2022), multiple recording sessions from the same subjects were treated as individual data points.

### Correlation of *GS* and mean DFA

Pearson’s linear correlation analysis was used to estimate correlation between subjects/sets’ *GS* and mean DFA values for each frequency. To obtain surrogate distribution of correlation cofficients, the order of the dependent variables was shuffled 1000 times. Multiple hypothesis testing was corrected with the Benjamini-Hochberg method, by pooling together both connectivity metrics and all frequencies. We also estimated correlations between *GS* and mean DFA with a partial-quadratic model, i.e., a purely quadratic correlation where the linear trend in the dependent variables was partialed out. To test whether the relationship between synchrony and criticality was concave and not convex (peaks rather than dipping, with the concave inverted-U curve opening down), in addition to obtaining the R^2^ statistic, the coefficient of the quadratic term was multiplied with the sign (note that the y-axis is reversed in the main text subject correlations figure panels, so that the negative values indicating concave correlations are on top).

### Correlation NS and mean DFA

We next calculated linear and partial-quadratic correlations across subjects for each cortical parcel and frequency. In MEG data, every local *NS* value of the 400-parcel Schaefer atlas was correlated with the corresponding parcel DFA exponent across sets, with outlier rejection, surrogate calculation, and FDR correction (this time including also the 400 parcels) same as above. In SEEG data, contacts from all subjects were pooled to parcels of the 100-parcel Schaefer atlas, and the correlations between parcel *NS* values and DFA exponents were computed for all parcels containing at least 5 electrodes (after outlier rejection, resulting in 77/100 parcels). Linear and partial-quadratic correlations were then computed for these parcels in the same way as described for the subject level. For visualization purposes, we grouped frequencies in data-driven bands, individuated as the optimal community structure determined by the Louvain method (Blondel et al., 2008) of the self-similarity frequency-by-frequency matrix of the linear parcel correlations (delta, δ: 2–3 Hz; theta, θ: 4–7 Hz; alpha, α: 9–12 Hz; beta, β: 15–29 Hz; low-gamma lγ 40–65 Hz; high-gamma, hγ: 77–135 Hz; ripple-gamma, rγ: 165–225 Hz; *Supplementary Figure 5*). Results were similar for single frequencies not grouped into bands (*Suppl.Figure 6*) and if statistics were averaged into bands after correlations instead of before.

