## Supplemental Figures for "Brain criticality predicts individual synchronization levels in humans"

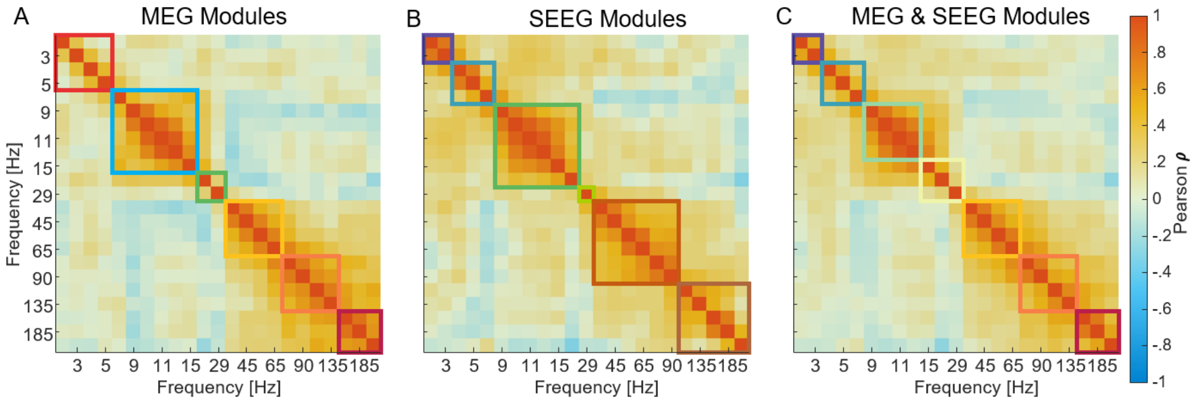


**Supplementary Figure 1.**

**A.** Spatial similarity analysis across frequencies using Louvain clustering (Blondel et al., 2008) for correlations of NS-DFA exponents (main text Figure 3A-B) in MEG. These Pearson coefficients were used as a directed weighted matrix in a multi-iterative Louvain community detection algorithm with the resolution parameter γ = 1.5 to identify modules (note that 0.5 ≤ γ ≤ 1 leads to only 2 modules, dividing low and high frequency bands at 30/40 Hz). **B**. Same as in A. but for SEEG contacts. **C** Same for combined MEG and SEEG data.

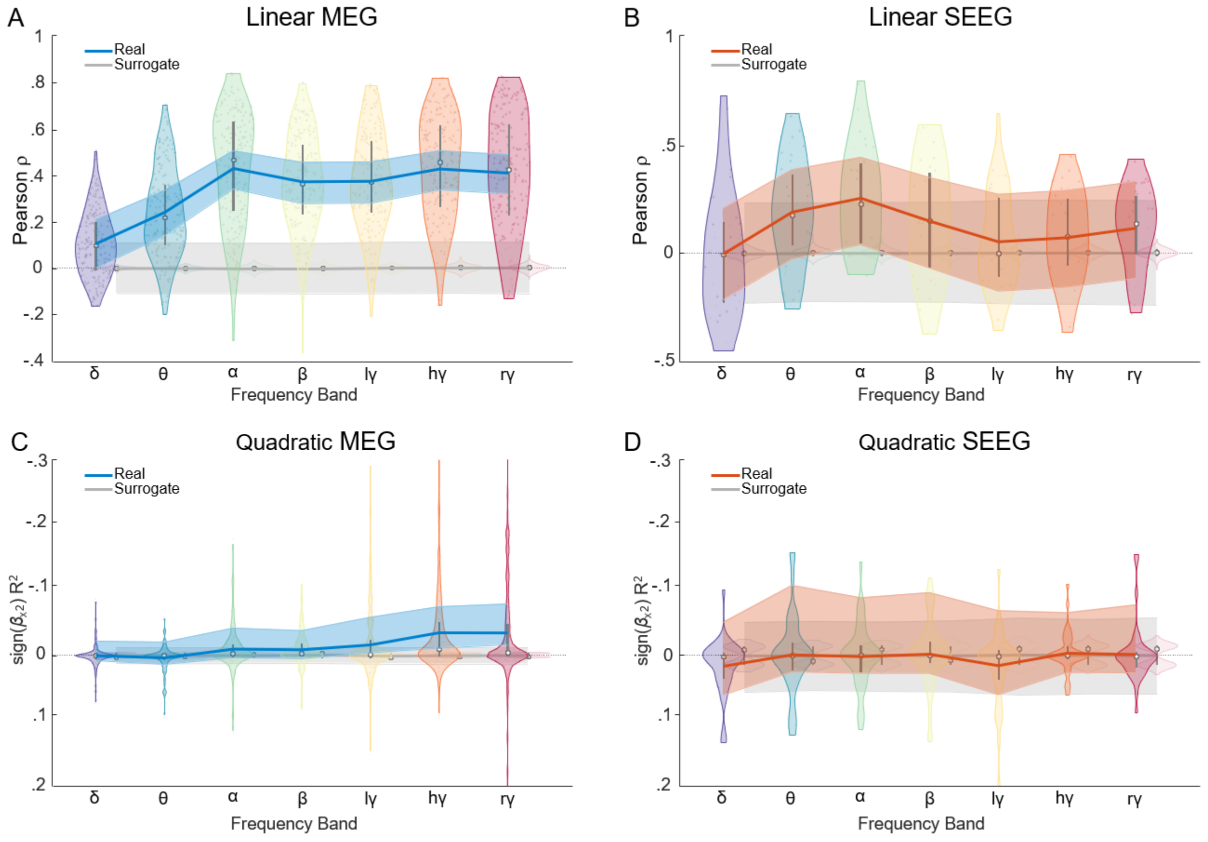


**Supplementary Figure 2.**

Linear and quadratic correlations of *NS* (MEG parcels and SEEG contacts) and DFA exponents. **A.** Means and distributions of linear Pearson correlation coefficients for different frequency bands in MEG data. Means of the real correlations are in blue, with shaded areas indicating 95% confidence intervals. Surrogate mean correlations obtained by case-resampling in grey, with the shaded areas indicating the 2.5-97.5$^{th}$ percentiles of the surrogate coefficients distribution. Violin plots show correlation coefficients distributions, with the median indicated by square and quartiles by notch indicators. **B.** Same as in *A.* for SEEG contacts. **C-D.** Same as above for partial quadratic (linear component removed) correlations multiplied with the sign of the quadratic coefficient (notice the y-axis has negative values on top).

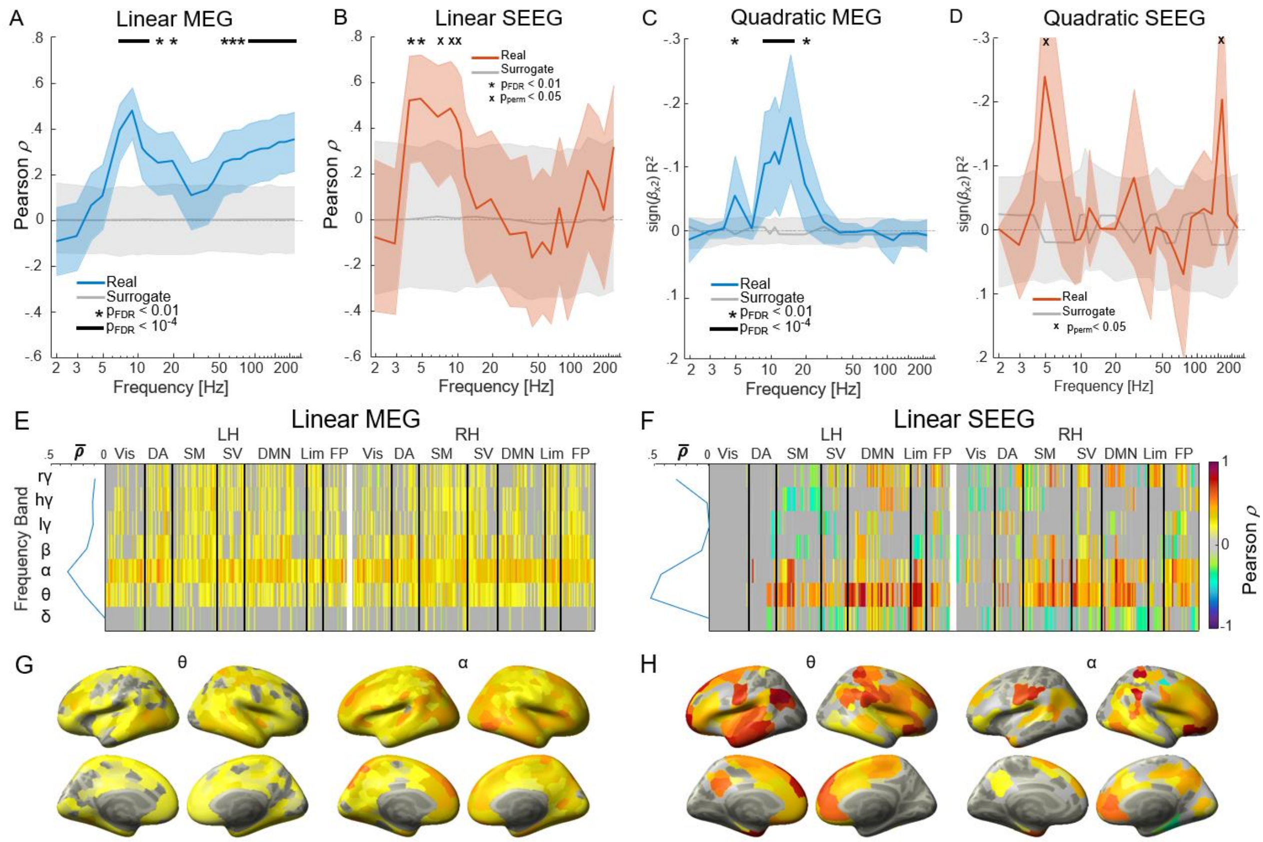


**Supplementary Figure 3.**

**A.** Linear between-subjects correlations of GS and mean DFA using PLV for MEG and **B**. wPLI for SEEG (the opposite connectivity metrics as were used in main figure). Correlations with 95% confidence intervals in blue or red, respectively, with grey indicating 2.5-97.5th percentiles surrogate distribution. Asterisks at the top indicate $p_{FDR}$ < 0.01 and the black line $p_{FDR}$ < $10^{-4}$. **C.** Partial quadratic correlations for of MEG and. **D.** for SEEG. **E.** The average correlation values in brain across parcels for MEG and **F.** for SEEG. **G.** Cortical correlations topographies of MEG *NS* and DFA for the theta and alpha frequency bands. **H.** Same as in *G.* for SEEG.

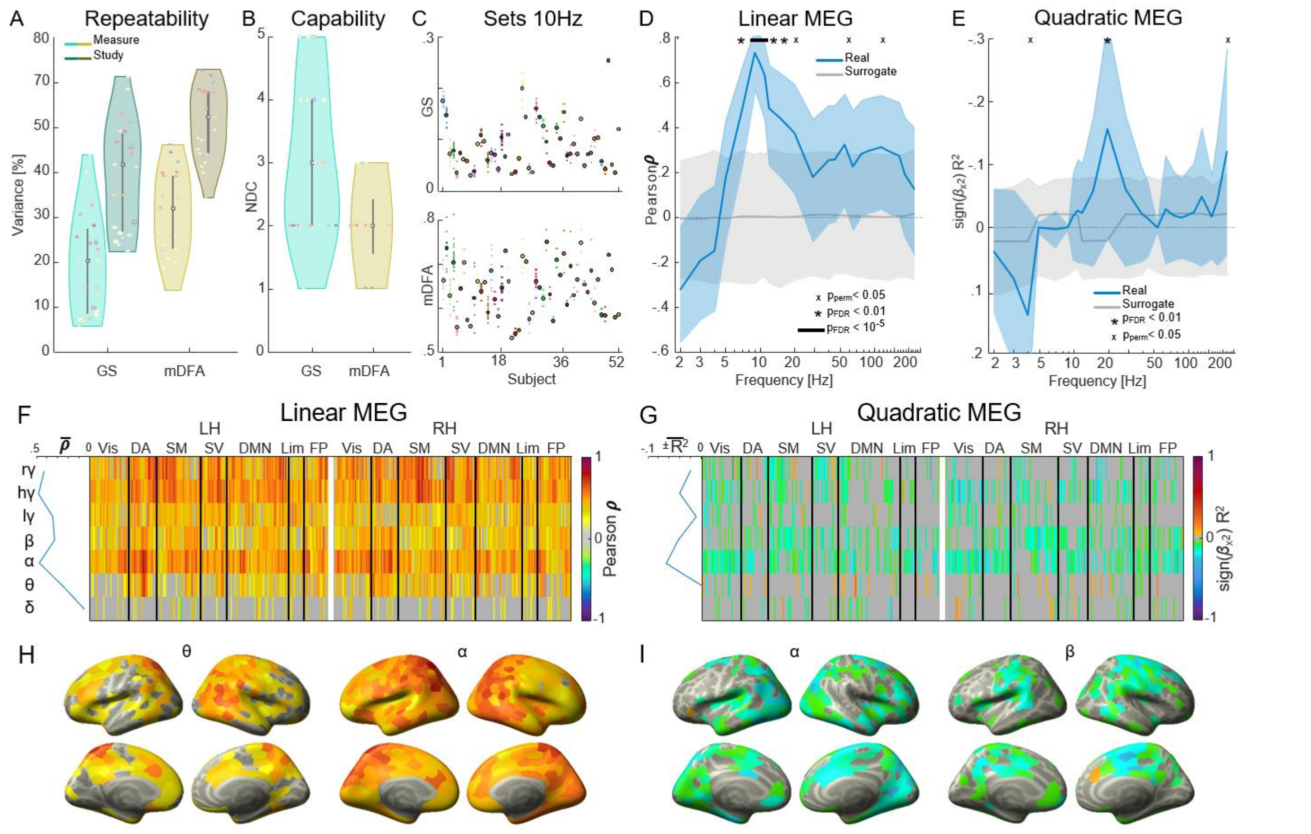


**Supplementary Figure 4.**

Repeatability of measures. **A**. Gauge Repeatability (ANOVA, Burdick et al., 2005) of MEG wPLI GS and mDFA of individual sessions. Violin plots show repeatability distributions of colour-coded frequencies as percentage of contributing variance, with median and quartiles notches. Darker violin plots show the study-wide variance. **B**. Single frequencies distributions of the capability of the metrics as Number of Distinct Categories (NDC). Larger points in the density violins are for frequencies with NDC > 3. **C**. Example MEG GS (top) and mDFA (bottom) for 10 Hz across session for each subject. The subject means, indicated with a bigger circle, were used in the subsequent analyses of this figure. **D**. Linear Pearson correlation coefficients of subject-averaged MEG GS vs. mDFA across frequencies with the 2.5-97.5th percentiles surrogates in grey. Xs indicate values beyond the 95% distribution of the surrogate correlations; asterisks at the top indicate $p_{FDR} < 0.01$; the black line $p_{FDR} < 10-4$. **E**. Partial quadratic correlationsof subject average MEG GS vs. mDFA with the R2 and the sign of the quadratic beta. **F**. Left and right hemisphere parcel-by-frequency-band matrices of linear correlations of subject-averaged MEG NS and DFA exponents (like Figure 3). **G**. Same as in F. for partial quadratic correlations times the sign of the quadratic coefficient. **H**. Cortical correlations topographies of subject average MEG NS and DFA for the theta and alpha frequency bands. **I.** Same as in H. for partial quadratic with the quadratic sign for the alpha and beta frequency bands.

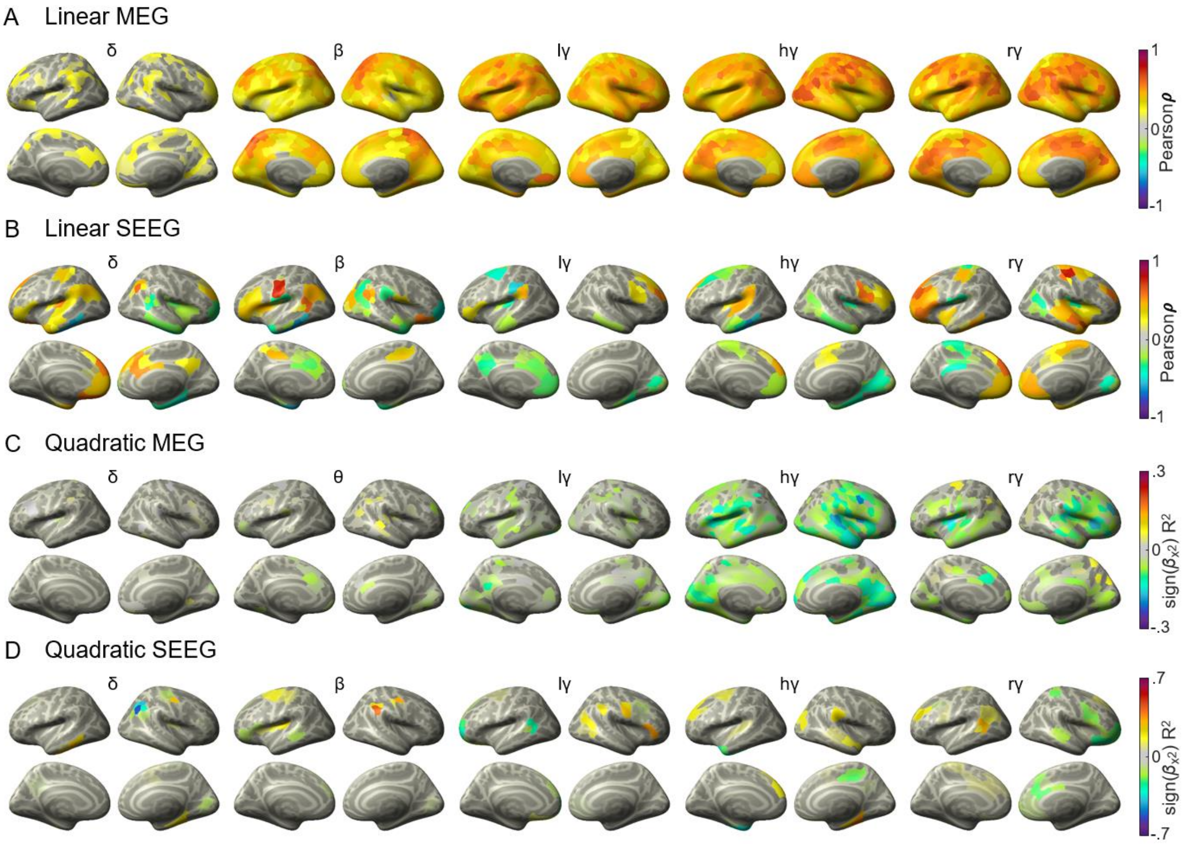


**Supplementary Figure 5.**

Cortical topographies of linear and quadratic between-subjects correlations of NS DFA for all the frequency bands not shown in Figure 3.

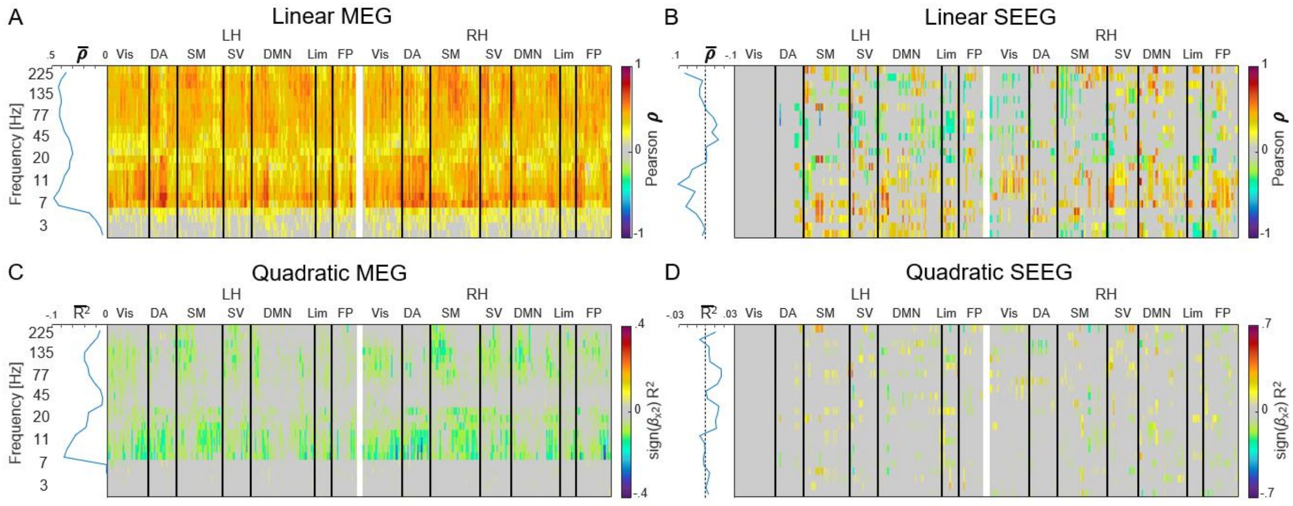


**Supplementary Figure 6.**

Linear and quadratic between-subjects correlations of local connectivity *NS* and DFA exponents for all parcels and single frequencies (not bands as in *Figure 3*). **A.** Left and right hemisphere parcel-by-frequency-matrices of linear correlations of MEG *NS* and DFA. Masked in grey are non-significant correlations. On the left side are the average correlation values of frequencies across significant parcels. **B.** Same as in *A.* for SEEG. **C-D.** Same as above for partial quadratic (linear component removed) with the sign of the quadratic coefficient.
